# PyamilySeq: Exposing the fragility of conventional gene (re)clustering and prokaryotic pangenomic inference methods

**DOI:** 10.1101/2025.05.30.657108

**Authors:** Nicholas J Dimonaco

## Abstract

Pangenomics has become a central framework for exploring microbial diversity and evolution, enabling researchers to distinguish genes that define shared biological function from those that drive adaptation. However, this relies on clustering genes by sequence similarity, a process that is far less deterministic than often assumed. This study introduces PyamilySeq, a transparent and flexible toolkit designed to diagnose and quantify hidden biases within gene clustering and pangenome inference methodologies. Using PyamilySeq, we can see how clustering thresholds (often hardcoded and poorly documented) and paralog handling can substantially alter gene family composition. Surprisingly, even parameters unrelated to clustering, such as decimal precision (0.8 vs. 0.80), output selection, and even CPU and memory allocation, can alter gene family assignments, challenging the assumption that identical clustering thresholds yield consistent results. Furthermore, tools often fail to report biologically meaningful or representative sequences for gene families, undermining downstream analyses. These findings reveal systematic fragilities in gene clustering and pangenome construction and highlight that pangenomics is not merely a data-driven task but a methodological one, where transparency, reproducibility, and interpretability are as critical as biological insight. This work calls for a re-evaluation of how pangenomes are constructed and compared, and advocates for methodologies that make their assumptions explicit and their results verifiable.

## 1 Introduction

Gene clustering, the process of grouping genes by shared characteristics such as function, structure, or, most often, sequence similarity, is essential for understanding microbial diversity and functional potential. As genome databases expand in both size and complexity, traditional deterministic approaches to sequence comparison are increasingly replaced by heuristic algorithms to improve computational efficiency. Yet these heuristic approaches introduce approximations that can compromise clustering accuracy [1]. Nevertheless, one key approach to derive biological significance from gene clusters is pangenome inference. Here, a pangenome represents the current set of proteincoding genes found within a species, including core genes present in all (or most, depending on the definition) genomes and varying levels of accessory genes that drive genetic diversity and adaptation [2]. Pangenomics play a crucial role in fields such as antimicrobial resistance surveillance, pathogen detection, and evolutionary analysis by identifying the sets of core and accessory genes that influence adaptation, virulence, and resistance across a species’ range [3]. However, the accuracy of these insights hinges on reliable genome assembly, annotation, and gene clustering, as errors in defining gene families can propagate through analyses and mislead biological interpretation [4, 5, 6]. Contemporary pangenome tools are complex software systems that integrate heuristic sequence similarity and clustering methods such as BLAST [7] and CD-HIT [8]. Despite their widespread use, these sequence clustering techniques and the pangenome tools that depend on them **often lack algorithmic transparency and flexibility**, raising concerns about their accuracy and reliability.

Pangenome tools typically apply additional post-clustering algorithms and grouping thresholds to refine clustering outcomes, yet the rationale behind critical steps, such as sequencing error handling, horizontal gene transfer detection, or paralog differentiation, is often poorly documented or difficult to interpret. This lack of clarity complicates efforts to determine whether observed gene clusters, or families, reflect true biological relationships or are artefacts of the tool’s methodology. While some studies have assessed the broad impact of biases introduced by gene clustering and pangenome inference methods [9, 10, 11], a systematic and detailed evaluation of their effects is lacking.

As mentioned, genome assembly and annotation remain major sources of variability, often introducing inconsistencies that directly influence clustering accuracy [12, 13]. The completeness of gene function classification remains a significant problem to be resolved for even the most well-studied genomes, such as *E. coli* which has only around 60% of its genes “Well-characterized” as recently as 2024 [14]. However, the impact of these factors on clustering results remains poorly understood. Although some tools attempt to correct annotation inaccuracies before clustering, they still rely on previous knowledge, alignment to known genes, or require additional experimental evidence, such as RNA and proteomic analysis [15, 16]. The extent to which annotation errors and clustering biases collectively distort pangenome inference is an open question that must be explored in greater depth.

A variety of prokaryotic gene-centric pangenome tools exist, each employing overlapping and unique methodologies. For example, PIRATE [17], PPanGGOLiN [18], PEPPAN [19], and PanTA [20] each provide their own approach to pangenome construction while employing many of the same heuristic methods for gene clustering (BLAST [7], DIAMOND [21], MMseqs2 [22], CD-HIT [8], etc). In this study, we focus on two representative tools: Roary [23], one of the older and more widely adopted tools with nearly five thousand citations to date, and Panaroo [16], which represents a more recent advancement in computational pangenome analysis. These tools not only exemplify the underlying techniques used by most pangenome software but are also among the most widely adopted and cited by the research community.

Roary [23], once a cornerstone of the field, has not been updated since 2019 (github.com/sanger-pathogens/Roary). The project’s GitHub repository currently states: *“PLEASE NOTE: we currently do not have the resources to provide support for Roary, so please do not expect a reply if you flag any issue”*. Consequently, the repository now primarily serves as a platform for community discussion, where users have shared unresolved issues, feature suggestions, and bug reports. Despite this, Roary remains widely used and is still incorporated into contemporary workflows [24]. Unfortunately, this situation is common in bioinformatics, where essential tools become unsupported over time due to the lack of funding priority and resources. In contrast, Panaroo [16] continues to be actively developed and was built with assembly and gene prediction error correction and improved gene relationship modelling.

A key factor influencing the accuracy of both Roary and Panaroo is their reliance on external clustering algorithms, particularly CD-HIT [8]. Designed for high-throughput sequence similarity clustering, CD-HIT plays a fundamental role in many bioinformatic workflows. However, its reliance on often misunderstood algorithmic decisions, one-size-fits-all parameters, and heuristic methods means that small parameter and input changes can drastically alter clustering outcomes. These rigid parameters, often set arbitrarily by both developer and user, and are sometimes based on historical defaults, may force unrelated genes into the same group or split genuinely related genes apart, leading to biased representations of genomic relationships. Understanding how these biases manifest across different tools and parameter settings is crucial for improving pangenome inference.

Given the complex interplay between annotation quality, sequence clustering, and pangenome inference, there is a pressing need for tools to systematically examine how methodological choices influence gene clustering and, therefore, gene family composition. To address this, PyamilySeq was developed as a diagnostic framework for studying both gene clustering and pangenome inference methodologies. However, throughout its development, PyamilySeq evolved into a versatile and fully-featured pangenome analysis tool. It not only replicates the core functionalities of widely used pangenome tools including Roary and Panaroo, but also their outputs and gene family designations as follows: Core genes, Soft core genes, Shell genes and Cloud genes. PyamilySeq also provides increased transparency in clustering decisions, flexibility in parameterisation, and iterative reclustering capabilities. In doing so, PyamilySeq serves both as a standalone pangenomic toolkit and an investigative engine for probing the assumptions and biases that underpin contemporary gene clustering methodologies. By varying key parameters and analysing their impact on gene family composition, we can see how clustering thresholds, algorithmic decisions, sequence alignment techniques, paralog handling and even runtime factors such as CPU (Central Processing Unit) and memory allocation shape pangenomic outputs. While PyamilySeq was developed with processes that may be broadly useful beyond this study, the objective is not to position it as a replacement for existing tools. Rather, this work aims to critically evaluate current methodologies, offering deeper insight into the factors shaping clustering results and guiding the development of more transparent, flexible, and biologically meaningful pangenome analysis approaches.

## 2 Methods

PyamilySeq is a pangenome analysis toolkit designed to provide transparency and user control at each stage of gene clustering and pangenome inference. An overview of how the main workflow of PyamilySeq operates is presented in Figure 1, and the complete list of software, including versions used, is available in Supplementary Section 4. It is implemented in Python and available as PyPi (https://pypi.org/project/PyamilySeq/) and bioconda (https://anaconda.org/bioconda/pyamilyseq) packages. The following sections describe the datasets and tools used to investigate the limitations of microbial gene-centric pangenomics.

**Figure 1:**
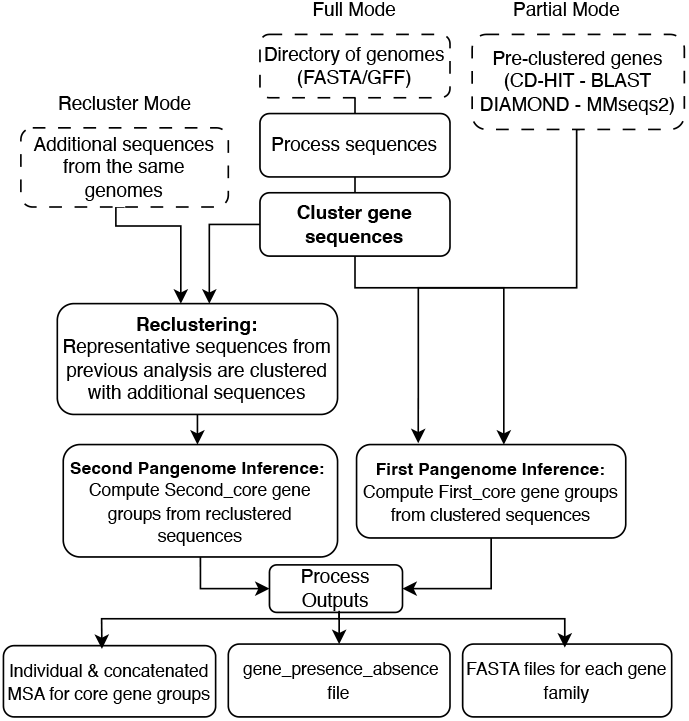
PyamilySeq toolkit overview. PyamilySeq offers two main operational modes: Full Mode, which processes genome annotations (GFFs and/or FASTA) and performs *de novo clustering* with CD-HIT, and Partial Mode, which allows users to input pre-clustered genes for pangenome inference. A third option, Recluster Mode, allows for additional sequences to be added to a previously performed pangenomic analysis. The toolkit outputs key files including multiple sequence alignments for core genes, gene presenceabsence matrices, and FASTA files for each gene group, providing a comprehensive framework for pangenome analysis and comparison.

### 2.1 Data selection and use

#### 2.1.1 10 *Escherichia coli* genome dataset

The first dataset used in this study was a set of 10 *E. coli* genomes, which varied widely in genome fragmentation (1 to 412 contigs) and in the number of protein coding genes (4,540 to 5,565). This collection of genomes was selected and downloaded from release 59 of Ensembl Bacteria [25]. To verify taxonomic assignments, knowing that incorrect assignments can lead to underestimation in pangenomic inference, the genomes were re-evaluated as *E. coli* using GTDB-TK [26]. This ‘diversity’ of assembly quality and gene number allowed for an assessment of PyamilySeq’s performance and utility on a small dataset with varying levels of genome and annotation ‘completeness’ (see Supplementary Table 1 for more details).

For sequence clustering, CD-HIT was used with its default parameters, including a sequence identity threshold of **0.90** but a modified `length difference cutoff` of **0.80**. Analysis was performed on both the DNA and AA sequences and is reported separately (cd-hit-est for DNA and cd-hit for AA). In parallel, pangenome analysis was performed on the same dataset using both Roary and Panaroo. Please see Subsection 2.3 for information regarding runtime parameters.

#### 2.1.2 74 *Escherichia coli* genome dataset

To demonstrate a more representative use case for a pangenome study, a larger collection of *E. coli* genomes were downloaded from Ensembl Bacteria (release 59). In an attempt to select for ‘highquality’ genomes, those with more than five fragments (excluding plasmids) were filtered out. This aimed to reduce the number of genomes with potential assembly and thus annotation errors. Using the same GTDB-TK taxonomic validation as before, three of the genomes initially labelled as *Escherichia coli* were identified as belonging to the genus *Citrobacter*. Following their removal, a final dataset of 74 genomes was collected and pangenomic inference was performed as detailed in Supplementary Table 2.

### 2.2 The PyamilySeq toolkit

PyamilySeq is a modular toolkit that replicates and dissects the key stages in typical pangenome workflows, including gene clustering, core/accessory gene classification, and alignment. This design enables users to examine how parameter choices and algorithmic behaviour influence outcomes at each step, providing transparency into the assumptions, limitations, and variability inherent in widely used pangenomic approaches.

Broadly, it begins by parsing the output of clustering tools such as CD-HIT, traces the origin of each gene within each cluster back to their source genome, and determines the distribution of each gene family e.g., core, soft-core, or accessory, across the dataset. To accommodate different definitions of gene family categories, PyamilySeq reports each classification using clear and user-definable percentage ranges for easier comparison, allowing users to align classifications with their dataset’s biological distribution. The default pangenome metrics reported by PyamilySeq (Core genes (99% `≥` strains = 100%), Soft core genes (95% `≥` strains *<* 99%), Shell genes (15% `≥` strains *<* 95%) and Cloud genes (0% `≥` strains *<* 15%))) align with the definitions of both Roary and Panaroo. To identify that these metrics are from a ‘first’ round of clustering (comparable to other pangenome inference tools - see Section 2.2.3 for details on reclustering), they are reported as: **First_core_99, First_core_95, First_core_15**, and **First_core_0**.

PyamilySeq is particularly suited for identifying subtle differences introduced by small parameter changes or input variations, making it valuable for sensitivity analyses and method comparisons. It also includes a suite of auxiliary tools for interrogating intermediate files and interpreting both inputs and outputs across runs of PyamilySeq and other pangenome tools. Users can perform initial clustering externally with their method of choice, including CD-HIT, BLAST, DIAMOND, or MMseqs2, and import results for pangenomic analysis. This is achieved through support for CD-HIT’s .clstr format and any tab-separated file formatted as *Node1* `\`*t Node2* (such as BLAST’s outfmt 6), representing relationships between gene sequences or clusters. To ensure interoperability, Seq-Combiner has been built to prepare a directory of GFF/FASTA files with the appropriate sequence IDs ready for clustering (see Supplementary Subsection 5.1). This flexibility, combined with the ability to handle both DNA and amino acid sequences as well as non-coding genes, allows researchers to tailor analyses to various genomic contexts.

Outputs include a *summary statistics*.*txt* file following a format similar to Roary and Panaroo, with additional modifications reflecting extended functionality (see Supplementary Listing 1), a presence-absence file matching Roary’s format for compatibility with existing downstream tools, individual and combined FASTA files containing gene sequences for each identified gene family, and a core gene alignment using MAFFT [27]. This level of customisation ensures users maintain control over the clustering process, enabling more accurate and meaningful results tailored to specific research needs.

PyamilySeq operates in two primary modes: **Full Mode** and **Partial Mode** (see Supplementary Figure 1). The following sections describe the main tools along with the data formats they take or create, all of which are available on GitHub (https://github.com/NickJD/PyamilySeq) and as part of the unified PyamilySeq PyPi package (https://pypi.org/project/PyamilySeq/).

#### PyamilySeq: Full Mode

Full Mode in PyamilySeq provides a complete gene clustering and pangenome analysis pipeline, automating the processing of annotated genomes to generate gene families and pangenome statistics. This mode is designed for users starting with annotated GFF or FASTA files and produces a gene presence-absence matrix by default. The workflow broadly consists of the following steps:

1. **Input:** PyamilySeq reads genomic data from a directory, accepting combined GFF+FASTA files, separate but linked GFF and FASTA files or FASTA files containing extracted gene sequences for each genome as input. Users can specify which genomic features to extract (e.g., CDS, rRNA, pseudogenes), ensuring compatibility with various annotation formats.
2. **Clustering:** Genes are clustered into families using CD-HIT, with user-definable thresholds for sequence identity and length difference. Clustering can be performed in either nucleotide or amino acid space.
3. **Pangenome Inference:** Gene families are classified as core, soft-core, or accessory genes based on customisable presence thresholds across genomes.
4. **Gene Presence-Absence Matrix:** PyamilySeq generates a matrix showing the distribution of gene families across genomes, formatted for downstream visualisation and comparative analyses.
5. **Optional Core Genome Alignment:** Representative core genes can be aligned using MAFFT, enabling concatenated sequence generation for phylogenetic studies.

Full Mode is ideal for researchers seeking an end-to-end solution for pangenome analysis, providing transparent gene clustering, classification, and phylogenetic insights directly from annotated genome data.

#### 2.2.2 PyamilySeq: Partial Mode

Partial Mode provides greater flexibility by allowing users to input pre-clustered gene sets instead of running CD-HIT clustering within PyamilySeq. This mode is designed for cases where external tools, custom clustering methods, or specific parameters have already been applied, and users want PyamilySeq to perform the final stages of pangenome inference and output generation. The workflow consists of:

1. **Input:** Users provide pre-clustered gene sets in either CD-HIT format (.clstr file) or as an edge-list file (`Node1 \t Node2`), ensuring compatibility with alternative clustering methods such as MMseqs2, DIAMOND, or BLAST.
2. **Pangenome Inference:** PyamilySeq processes the pre-clustered data to classify genes into core, soft-core, and accessory categories, applying the same customisable presence thresholds as available in Full Mode.
3. **Gene Presence-Absence Matrix:** A presence-absence matrix is generated from the provided clusters, formatted for downstream visualisation and comparative analysis.
4. **Optional Core Genome Alignment:** If selected, PyamilySeq aligns representative core genes using MAFFT, following the same approach as Full Mode.

#### 2.2.3 PyamilySeq: Reclustering

To support ongoing genome annotation research, PyamilySeq includes a **reclustering** feature that allows newly annotated gene sequences to be integrated into existing pangenome results and gene clusters. This is especially useful when additional gene sequences become available after initial clustering and allows users to **update and refine pangenome inferences without reprocessing entire datasets and losing the initial gene family assignments**. We are able to follow the positioning of the new sequences in the context of the original clustering, determining whether they fit within established clusters or if they form entirely new ones. This approach not only improves the accuracy of gene family assignment, but also reveals previously undetected genetic elements that may have been overlooked in the initial analysis.

A key example of this approach can be seen in the StORF-Reporter study [28], which demonstrated how the integration of StORFs (Stop-Open Reading Frames), often longer than standard gene annotations, had a significant impact on gene clustering. Due in part to their often-extended lengths, StORFs had the potential to either **merge or fragment** existing gene families. In that study, accurately reporting the true distribution of these newly annotated sequences required a **reclustering approach**, a feature absent in other tools. Panels **A** and **B** in Figure 2 illustrate how the addition of longer sequences can dramatically alter clustering results, especially in length-first clustering approaches such as CD-HIT.

**Figure 2:**
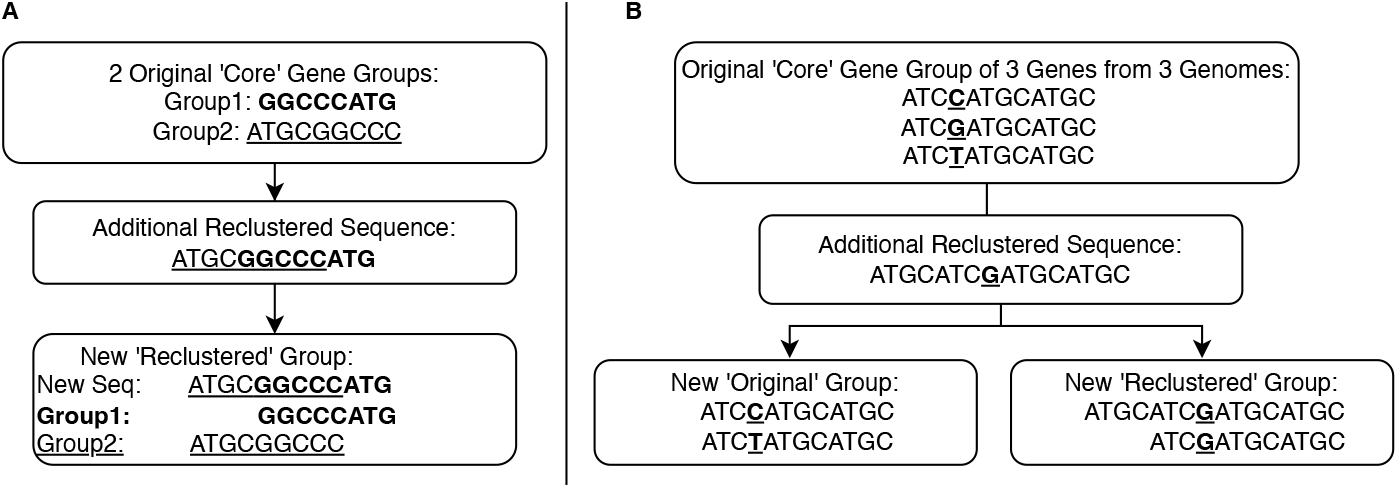
Reclustering additional sequences can fragment original clusters, decreasing the number of core/soft-core gene groups.

To track both existing and additional sequences during PyamilySeq’s reclustering step, additional metrics were required. The four additional categories reported are: **extended** (gene families moved to higher prevalence categories by the addition of new sequences), **combined** (gene families created by merging two or more original families through the addition of new sequences), **Second** (gene families defined exclusively from their constituent new sequences), and **only Second** (Second category families containing no original sequences from first-round clustering). Further details are provided in Supplementary Listing 1.

### 2.3 Parameter selection

Unless otherwise specified, the pangenome tools in this study were used with the following parameters.

#### Roary v3.13.0 parameters

All parameters were set at defaults apart from: CPU allocation at 8 `(-p 8)` and create a multifasta alignment with MAFFT `(-e -n)`.

#### Panaroo v1.5.1 parameters

All parameters were set at defaults apart from: CPU allocation at 8 **(-t 8)**, stringency mode of moderate `(--clean-mode moderate)` and create a core gene alignment `(-a core)`.

#### PyamilySeq v1.2.0 parameters

All parameters were set at defaults apart from: CPU allocation at 8 `(-T 8)`, Memory allocation of 4,000MBs `(-M 4000)`, a CD-HIT `identity threshold` of 90% `(-c 0.90)`, a `length difference cutoff` of `0.80` and create a core gene alignment `(-a core)`. Additionally, PyamilySeq has been designed to expand all single decimal place parameters to two decimal places `(0.0 to 0.00)`.

### 2.4 Robinson-Foulds analysis of inferred phylogenetic trees

To further confirm the similarity of the phylogenetic trees generated by each tool and dataset, Robinson-Foulds (RF) [29] symmetric difference tests are performed on the trees produced by FastTree2 [30] from the concatenated core gene MSA’s. This is done using the Python package ETE 3 [31]. RF can be used to measure the dissimilarity between two phylogenetic trees by quantifying the number of shared splits and normalising it to a maximum possible distance, typically ranging from 0 to 1, where 1 is the most different.

## 3 Results

Roary, Panaroo and PyamilySeq are applied to two datasets: one comprising 10 *E. coli* genomes with varying levels of assembly fragmentation, used to examine how assembly quality may influence pangenome inference; and a second dataset of 74 *E. coli* genomes, each exhibiting fewer than five contigs, providing a high-quality, ‘well-characterised’ genome set for robust comparative analysis. PyamilySeq can perform analysis of gene families using either nucleotide (**DNA**) and amino acid (**AA**) sequences. This dual approach lets us explore limitations and differences inherent to clustering and alignment tools operating in DNA or protein space. Throughout this study, **PyamilySeq DNA** and **PyamilySeq AA** will be used to explicitly state which sequence type analysis is being performed or discussed. An overview of how PyamilySeq works is presented in Figure 1.

### 3.1 Pangenomic analysis of 10 *Escherichia coli* genomes reports a consensus of approaches

First, PyamilySeq was applied to the set of 10 *Escherichia coli* genomes with relatively a high sequence identity fixed to 0.90 and a `length difference cutoff` of 0.80. From the results summarised in Table 1, we can make a number of key observations. Notably, the number of core gene groups reported by PyamilySeq falls just slightly below the results reported by Roary and Panaroo. This initial comparison serves as a valuable ‘sense check’, demonstrating that CDHIT alone (with no post-clustering algorithms and when configured with relatively strict clustering parameters), through PyamilySeq’s inference, aligns reasonably well with established tools. Such results suggest that PyamilySeq can provide a reliable basis for exploring the influence of clustering parameters on gene family assignments, helping to identify potential biases and opportunities for refinement in clustering and pangenomic methods. The phylogenetic trees inferred from each of the three tools are presented in Supplementary Figure 2. The lack in structural differences between the trees is backed up by normalised RF distances of 0, demonstrating that they yield consistent results when analysing this relatively small but fragmented set of genomes.

**Table 1:** Table reporting the gene family composition for each tool with default parameters on the two sets of 10 and 74 *E. coli* genomes.

| Groups | 10 *E. coli* genomes |  |  |  | 74 *E. coli* genomes |  |  |  |
| --- | --- | --- | --- | --- | --- | --- | --- | --- |
|  | PyamilySeq[DNA] | PyamilySeq[AA] | Roary | Panaroo | PyamilySeq[DNA] | PyamilySeq[AA] | Roary | Panaroo |
| Core genes | 2,989 | 2,996 | 3,078 | 3,309 | 432 | 357 | 1,023 | 2,824 |
| Soft core genes | 0 | 0 | 0 | 0 | 2,121 | 2,139 | 860 | 472 |
| Shell genes | 3,309 | 3,259 | 3,463 | 3,027 | 3,185 | 3,220 | 4,696 | 3,040 |
| Cloud genes | 5,314 | 5,469 | 6,181 | 2,607 | 34,908 | 37,014 | 31,890 | 15,190 |
| Total genes | 11,612 | 11,724 | 12,722 | 8,943 | 40,646 | 42,730 | 38,469 | 21,526 |

### 3.2 Pangenomic analysis of 74 *Escherichia coli* genomes reports significant differences based on approach

Although the previous analysis of 10 *E. coli* genomes showed minimal differences between clustering approaches; most pangenomic studies involve a larger and more diverse set of genomes. Expanding the analysis to 74 *E. coli* genomes reveal substantial discrepancies in core and accessory gene assignments, highlighting the impact of clustering methodology on pangenome inference. The results are summarised in Table 1.

As the number of *E. coli* genomes analysed increases, discrepancies between the three tools become more pronounced as each tool now reports a substantially different number of core genes. Using DNA-based clustering, PyamilySeq identifies 432 core gene families, compared to 357 using its AA-based clustering, both significantly lower than the 1,023 core genes reported by Roary. Panaroo, in contrast, reports 2,824 core genes, nearly three times as much as Roary and almost seven times more than PyamilySeq (Table 1). These discrepancies raise some fundamental questions: Which tool provides the most accurate representation of ‘this’ pangenome? Or, do these differences instead reflect the inherent complexity and subjectivity of pangenome analysis itself? Somewhat surprisingly, despite these substantial differences in core gene counts, the phylogenetic trees built from each tool’s core gene families (Supplementary Figure 3) appear broadly similar. The RF analysis further supports this, with the RF distances between the phylogenetic trees being as follows: the comparison between Roary and PyamilySeq reported a normalised distance of 0.18310; Roary and Panaroo reported a normalised distance of 0.15493; and the comparison between Panaroo and PyamilySeq reported a normalised distance of 0.16901.

Although this analysis focuses on a single species, it may explain why discrepancies in core gene counts often go unreported in other studies. If the primary objective of a pangenome study is phylogenetic inference, and core gene count variations do not substantially alter tree topology, such discrepancies may remain undetected or be considered inconsequential. However, this does not mean such discrepancies are scientifically acceptable. While phylogenetic reconstructions may tolerate some variation in gene family assignments, functional analyses, such as studies examining metabolic capacity, virulence potential, or evolutionary conservation, are inherently more sensitive to how core and accessory genes are defined.

To investigate whether core gene variation affects functional interpretation, we can examine the functional annotations (assigned by eggNOGmapper [32]) of genes classified as core under different methods. Under a straightforward hypothesis, that if a tool inflates its core gene collection by misclassifying accessory genes as core, we could expect an increase in ‘POORLY CHARACTERIZED’ genes within the core set, as accessory genes are typically less studied and functionally annotated than ‘true’ core genes [**?**]. Our results, however, reveal the opposite pattern. Rather than observing an increase in ‘POORLY CHARACTERIZED’ genes as core size expanded, we found a proportional 6% increase in genes assigned to ‘METABOLISM’ categories alongside an actual decrease in ‘POORLY CHARACTERIZED’ genes. A chi-square test further positioned this shift as statistically significant (*χ*^2^ = 31.5353, *p <* 0.001). This result has important implications: core gene inflation does not simply misclassify a ‘random’ subset of accessory genes; rather, it looks to be recruiting specific functional categories into the core, fundamentally changing the perceived functional landscape of the species. Such systematic biases could lead researchers to draw incorrect conclusions about metabolic capabilities, evolutionary constraints, or functional repertoire, particularly when comparing pangenomes across studies that employ different tools or parameters. Supplementary Figure 4 and Supplementary Table 3 present broad functional classifications of the core gene families identified by each tool, assigned using eggNOG-mapper [32].

### 3.3 Influence of CPU, memory, parameter precision and core gene alignment on gene clustering results

During PyamilySeq’s development and testing, discrepancies were observed in gene grouping counts between Full and Partial modes, despite using identical CD-HIT clustering parameters. While it is well known that heuristic-based sequence alignment tools such as CD-HIT and BLAST can yield different results, the impact of system resource allocation, such as CPU thread count (selectable with ‘`-T`’) and memory provisioning (selectable with ‘`-M`’) on clustering results is less widely recognised. As seen in Supplementary Table 4, passing varying CPU and memory allocations to CD-HIT results in small but observable differences in the core gene assignments. These results suggest that computational resource allocation can influence clustering outcomes beyond just performance.

While the number of core genes remains almost unchanged across different CD-HIT parameter runs, some genes were reassigned to different clusters, indicating a rearrangement in their grouping. For example, in PyamilySeq’s default CD-HIT run (`-T 8 -M 4000`), the singleton gene ‘*ENSB nhmgO-nwCXecCuN*’ remained unclustered. However, under CD-HIT’s default settings (`-T 1 -M 800`), it clustered with another sequence, ‘*ENSB EF9iv1mvUVBoPlR*’ (PID: 90.16%), from a different genome. These results are consistent across multiple interactions with the same parameters, thus further indicating that they are a direct result of parameter changes and not just run-to-run heuristics. Notably, in the PyamilySeq (`-T 8 -M 4000`) run, ‘*ENSB EF9iv1mvUVBoPlR*’ was grouped with 25 other sequences across 22 genomes. To determine whether clustering discrepancies increase with higher CPU and memory settings, CD-HIT was executed with even higher values `(-T 12 -M 40000`). Surprisingly, ‘*ENSB nhmgO-nwCXecCuN*’ clustered with ‘*ENSB EF9iv1mvUVBoPlR*’, mirroring the default (-T 1 -M 800) results rather than the PyamilySeq ‘`-T 8 -M 4000`’ parameter run. While this is an issue specifically attributable to the CD-HIT clustering and not PyamilySeq (or Roary/Panaroo), it must be accounted for by at least reporting the performance parameters with the results of the analysis. Supplementary Table 5 reports the CD-HIT initialisation outputs for the aforementioned different `-T (CPU)` and `-M (Memory)` parameter choices. What we can see is CD-HIT utilising different numbers of ‘representatives’ and ‘word counting entries’ depending on the selection of CPU and Memory parameters, effectively creating unintentional seed changes (see Supplementary Listings 1-3 for a more detailed view).

Given that CD-HIT serves as the primary clustering step in both Roary and Panaroo, it was tested whether CPU allocation affected their gene family assignments on the set of 74 genomes. As shown in Table 2, even with this relatively small dataset, varying CPU allocations (2, 4, and 8) resulted in differences in the reported gene families for both Roary and Panaroo. As a consequence of this, all analysis reported for PyamilySeq, Roary and Panaroo use 8 CPU threads, and all runs of CD-HIT (internal or external to PyamilySeq) were allocated 4,000MB of memory.

**Table 2:** This table reports the different gene family results reported by Roary and Panaroo on the 74 **E. coli** dataset with default parameters but different CPU allocations.

| Groups | Roary 2 | Roary 4 | Roary 8 | Panaroo 2 | Panaroo 4 | Panaroo 8 |
| --- | --- | --- | --- | --- | --- | --- |
| Core genes | 1,030 | 1,031 | 1,023 | 2,824 | 2,824 | 2,824 |
| Soft core genes | 859 | 860 | 872 | 472 | 472 | 472 |
| Shell genes | 4,743 | 4,720 | 4,740 | 3,027 | 3,033 | 3,040 |
| Cloud genes | 31,759 | 31,824 | 31,709 | 15,195 | 15,163 | 15,189 |
| Total genes | 38,391 | 38,435 | 38,344 | 21,518 | 21,492 | 21,525 |

Another unexpected factor influencing CDHIT clustering results was the precision of the (`-c sequence identity threshold`) and ( `-s length difference cutoff`) parameters. Specifically, using one vs. two decimal places (e.g., `0.9` vs. `0.90`) led to differences in clustering outcomes. In the 74 genome *E. coli* dataset, these small variations resulted in some accessory genes being grouped differently. This effect possibly stems from differences in how programming languages handle floating-point precision and rounding. While detailing further as to why this happens is not within the scope of this study, the results are consistent across runs on both x86 Linux and ARM Apple Silicon systems. These findings suggest that the interaction between CPU allocation, memory, and parameter precision introduces a degree of non-determinism in clustering outputs. Complicating matters further, it remains unclear how Roary and Panaroo internally handle and pass user-defined parameters, making it difficult to fully control for these sources of variability. Perhaps the most unexpected clustering inconsistency arises when running Roary in alignment mode. As seen in Supplementary Table 6, enabling core gene alignment with either MAFFT or PRANK [33] influences the distribution of core and accessory genes. While this is an effect that may become more pronounced in larger datasets, even in this relatively small set of 74 genomes, enabling MAFFT alignment increases the core gene count from 1,2023 to 1,026 and decreases the soft core from 872 to 848, with difficult to determine inter-gene family rearrangements.

### 3.4 Does it matter whether we cluster on DNA or AA?

Gene clustering and thus pangenomic inference are most often performed using amino acid (AA) sequences, under the assumption that functional conservation is more accurately captured at the protein level. While considerable progress has been made in identifying protein-coding regions in microbial genomes [13], accurately predicting the precise amino acid sequence from nucleotide data remains challenging. Current computational methods still struggle with ambiguous start sites, non-standard codon usage, and frameshifts; issues that are especially problematic in diverse microbial communities [34]. Although AA-based clustering is the standard, it risks overlooking subtle but biologically meaningful variation present at the DNA level. For example, synonymous substitutions and regulatory sequence variation, which are invisible in the AA sequence, may reflect important evolutionary signals. As shown in Supplementary Table 7, sequences that are identical at the protein level can still exhibit nucleotide-level divergence, which may be informative for phylogenetic reconstruction or strain-level resolution. This raises the question of whether clustering on AA sequences alone is always appropriate, especially in studies that aim to capture fine-scale evolutionary or functional differences.

To investigate the differences between DNA- and AA–based sequence clustering, PyamilySeq was applied to both the 10 and 74 genome studies using both their nucleotide and protein sequences. While the total number of clusters using PyamilySeq remained nearly identical in the 10-genome study (2,989 core gene families in DNA vs. 2,996 in AA), a larger discrepancy emerged in the 74-genome study, where core gene families decreased from 432 (DNA) to 357 (AA). These small-scale examples illustrate key differences between both DNA and AA sequence clustering. To explore this further, the largest cluster identified in the 74-genome study was examined. In the DNA clustering, CD-HIT (cd-hitest) formed a 506 gene cluster, selecting *ENSB s-GX UiVmBsWL9K* (291 nt) as its representative sequence. The most divergent sequence within this cluster, *ENSB Vyjslictzez3CNX* (249 nt), had a reported 98.39% sequence identity to the representative. These two sequences effectively define the range of sequence and length similarity within this cluster. However, in the AA sequence clustering, both sequences were reported as singletons, despite clustering under identical default parameters. This discrepancy stems from an inherent limitation of CD-HIT’s clustering algorithm: gaps do not contribute to sequence differences, meaning alignments with an arbitrary number of gaps can still report 100% identity. Despite these clustering differences, the pairwise RF distances between DNA and AA trees were minimal (0 at 80% `length difference cutoff` and 0.070423 at 0%), demonstrating how downstream analyses such as phylogenetics can mask important methodological effects on variation in pangenome composition.

Although we can not directly visualise CD-HIT alignments, we can try to infer its behaviour using BLAST and Clustal-Omega [35, 36] by observing how other coding-frame-unaware sequence aligners operate. As shown in Supplementary Figure 5, both `blastn` and `blastp` report alignments between *ENSB s-GX UiVmBsWL9K* and *ENSB Vyjslictzez3CNX* with a percentage identity above 90%, whereas Clustal-Omega does not. Further to this, while a 90% identity cut-off in Clustal-Omega would exclude these sequences, their DNA alignment score of 89.3% is higher than their AA alignment of 78.3%. This discrepancy could potentially explain why CD-HIT clusters them in DNA but not in AA. Nevertheless, the alignments reveal gaps that may be misinterpreted by alignment tools lacking coding-frame awareness, such as CD-HIT and BLAST. Notably, the only annotated protein domain information for these sequences corresponds to the aligned segments, which are classified as intrinsically disordered regions, possibly further complicating reliable alignment and clustering.

Lastly, an extreme example that further highlights the challenges of using sequence clustering tools is the case of *ENSB U6Q8FqhU eaDP5e* and *ENSB 8ANT4ngdQh30mf9*, which are two sequences of vastly different lengths, 1,119 and 2,091 nt, respectively. `blastn` yields “no significant similarity” (see Supplementary Figure 6, Panel **A**), yet `blastp` aligns them with ~90% identity over their shared region (first alignment in Panel **B**). Panels **B** and **D** illustrate a scenario closer to the behaviour of CD-HIT, where higher levels of sequence identity for AA lead to clustering but not for DNA. Although likely rare, this observation can be partially explained by codon optimisation and reassignment [37]. Nonetheless, they underscore a broader issue that sequence alignment and clustering tools used for pangenomic inference are often ill-equipped to fully capture biological complexity.

### 3.5 Paralogs, orthologs and gene duplication

Pangenome inference tools aim to classify gene families by distinguishing between orthologs, genes that arise from speciation events and typically retain similar functions, and paralogs, genes that result from gene duplication within a genome and may evolve new or redundant roles. Accurate separation of these genes is crucial, as misclassification can obscure evolutionary relationships, affect gene family counts, and impact downstream phylogenetic and functional analyses [38].

Both Roary and Panaroo perform ‘paralog splitting’ by default, and we will now examine how it impacts their resulting pangenomes. As shown in Table 3, turning off paralog splitting in Roary has a substantial impact, nearly tripling the number of core gene families (1,026 to 2,554). This underscores the importance of paralog splitting and thus the impact that any potential problems with how it is conducted could have on the final output. To investigate this substantial increase, the *gene presence absence*.*csv* files from both Roary runs were compared, one with paralog splitting and the other without on the set of 74 genomes. The differences were striking. Under default parameters, Roary’s largest core gene group contained 149 sequences, and 420 gene families had at least one genome with multiple gene sequences, yet only 4 of these were core gene families. However, with paralog splitting turned off, the largest core gene group now had 478 sequences, and of the 5,472 gene families containing at least one genome with multiple gene sequences, 2,053 were core. These findings highlight the significant impact of paralog splitting on Roary’s output. Notably, this process does not lead to a reduction of the core, which might be assumed given that the objective of paralog splitting is to split apart gene groups that were previously combined, but quite the opposite.

**Table 3:** Gene family distribution of the 74 *E. coli* genomes reported by Roary with and without paralog splitting.

| Groups | Roary with splitting | Roary without splitting |
| --- | --- | --- |
| Core genes | 1,026 | 2,554 |
| Soft core genes | 848 | 602 |
| Shell genes | 4,763 | 2,430 |
| Cloud genes | 31,714 | 16,000 |
| Total genes | 38,351 | 21,586 |

To explore how Roary assigns suspected paralogs, we examined the core gene family with the most sequences (group 21463) in the `-s dont split paralogs` output. This family contained sequences from all 74 genomes, totalling 478 genes, reporting that multiple copies existed within most genomes. Using PyamilySeq’s `Seq-Finder` tool, these genes were cross-referenced in the default Roary output, revealing that they had originally been distributed across 10 separate gene families, each containing 1 to 70 genes (see Supplementary File 1). This reassignment in part explains why Roary reports far fewer core gene families under default settings. To further investigate, the DNA and AA sequences of these 478 genes were retrieved and clustered with CD-HIT. The results showed that all 478 AA sequences were identical (84 AAs, 100% identity). At the DNA level, only 6 sequences differed, each by a single inconsistent nucleotide. Notably, these variants had other identical copies within the same genome, suggesting recent duplication events. These findings indicate that Roary’s default settings, as the tool description states, likely rely on gene synteny and neighbourhood context rather than on sequence variation alone. While this is a powerful algorithmic choice, it will be reliant on high-quality assemblies and annotation to determine each gene’s placement. Further analysis of core gene families containing suspected paralogs revealed significant length variation. For example, in group 9 the average gene length was 690 nucleotides, yet gene lengths ranged from 119 to 6,767 nucleotides. AA alignment of the longest and shortest sequences in this group with Clustal Omega showed almost no similarity, reporting a coverage of 1.9% and percentage identity of 19.7% (See Supplementary Figure 7). While individual sequences were more similar to others in the group, the lack of overlap between the longest and shortest sequences raises concerns about Roary’s default clustering criteria.

Supplementary Figure 8 illustrates an example from Panaroo (default parameters) where clustering sequences of varying lengths may be biologically informative. In core gene family group 21184, the genome ‘40363 C01’ contains a 962 AA sequence, while ‘39282 C02’ has three shorter sequences aligning independently across different regions of the longer sequence (start, middle, and end). BLAST analysis reveals that the longer sequence is a 100% identity match to a known *E. coli* pitrilysin protein (`NCBI Ref: WP 038338917.1`), while the three shorter sequences also match the same protein, albeit with varying identities (See Supplementary Figure 8). Another example can be found in group 21164, where 73 of the 74 genomes contain two sequences, while a single genome, o25b h4 gca, is represented by one gene of 2,292 nt, which encompasses the aligned regions of all other shorter sequences within that family. This structure is illustrated in Supplementary Figure 9, where the first sequence represents the long gene from the outlier genome, followed by two pairs of shorter sequences (699 nt and 1,611 nt) and another smaller pair of (246 and 144 nt). This could be an example of Panaroo recovering gene fragments resulting from pseudogenisation or assembly/annotation error. Nevertheless, Roary under default parameters separates these sequences into separate groups but does combine some when run with `-s dont split paralogs`. Although grouping genes with significant length variation may be appropriate in some cases, concealing these differences within the clustering process risks misleading conclusions.

Lastly, this behaviour is supplemented by Panaroo’s ‘refound’ system, which attempts to recover genes that may have been missed during annotation or disrupted by assembly artefacts, using BLAST-based similarity searches. For example, group 21514 consists of two Ensembl annotated gene sequences (‘*ENSB rCdIN-kBoySgj1Z*’ and ‘*ENSB QFZ19zgUaerM5rc*’) and 72 refound sequences, making this a single-copy core gene family. Expectedly, when Panaroo is run with its refound system disabled, these two sequences are reported together in a group of their own. While it is difficult to exactly infer how Panaroo is identifying these 72 additional sequences, one can assume from the paper that it is checking through BLAST if the annotated sequence is present and unannotated in the other genomes [16]. However, the refound system is not without its own quirks and is known to return partial or fragmented sequences, especially in cases of gene pseudogenisation or poor-quality assembly. As such, it may introduce misleading or inconsistent results into the final gene family assignments, particularly concerning users who are unaware of its influence, as these clusters are not always clearly distinguishable from those not formed from refound sequences. While some of these issues have been acknowledged and partially mitigated in recent updates, the refound system remains enabled by default (https://github.com/gtonkinhill/panaroo/issues/263). Ultimately, this highlights a broader issue of transparency in pangenomic inference tools. Key decisions made, such as whether to group fragmented genes and how to resolve suspected paralogs, are often not easily accessible or visible to the user. This lack of clarity can mask biologically significant patterns or errors, and may contribute to conflicting or irreproducible results across tools and studies.

### 3.6 The impact of sequence length on gene clustering

We have seen several examples where gene sequences of vastly different lengths were clustered together by either Roary or Panaroo. To investigate this further, PyamilySeq was rerun on the 74-genome dataset with CD-HIT’s `length difference cutoff` set to its default of 0%. To assess the impact of these relaxed length constraints, PyamilySeq’s Group-Summary tool was applied to the amino acid clustering results. The results reveal that CD-HIT’s default parameters allow for the clustering of sequences with extremely different lengths because it calculates similarity based on local overlapping regions rather than the full length of both sequences (CD-HIT provides parameters to change this). Consequently, sequences sharing only a conserved domain or small high-similarity segment (*>*= 90% identity) are grouped together in the initial clustering step, a fundamental issue that can propagate through entire pangenome inference workflows. A summary is provided in Table 4, with the full dataset available in Supplementary File 2. As can be seen in Table 5, a dramatic shift in core and soft core gene counts occurs when adjusting the `length difference cutoff` in CD-HIT. Specifically, lowering the length difference threshold allowed genes with greater size disparities to cluster together, resulting in a sharp increase in core gene families (for example, from 357 at 80% to 1,953 at 0% `length difference cutoff`).

**Table 4:** Summary of 10 gene clusters with a diversity of sequence lengths but high sequence identity from the CD-HIT results of clustering AA sequences at 90% identity and 0% length difference cutoff on the 74 *E. coli* genomes.

| ID | # Seqs | # Genomes | Avg Len | Len Range | Avg Ident' |
| --- | --- | --- | --- | --- | --- |
| 0 | 745 | 61 | 124.51 | 67-232 | 96.23 |
| 1 | 502 | 58 | 199.87 | 29-409 | 97.17 |
| 9 | 213 | 33 | 184.5 | 57-449 | 96.87 |
| 17 | 153 | 74 | 510.87 | 88-1070 | 98.28 |
| 19 | 152 | 74 | 618.18 | 84-1286 | 99.31 |
| 20 | 152 | 74 | 349.03 | 86-736 | 98.69 |
| 21 | 150 | 74 | 439.54 | 64-955 | 98.59 |
| 55 | 127 | 63 | 411.06 | 71-799 | 95.57 |
| 57 | 125 | 63 | 430.02 | 77-884 | 94.07 |
| 63 | 115 | 59 | 463.9 | 31-1016 | 96.37 |

**Table 5:** PyamilySeq gene group outputs of different CD-HIT length difference cutoff’s of 0%, 40% and 80%. Results for both DNA and AA clustering are shown. LD = Length Difference.

| Groups | DNA |  |  | AA |  |  |
| --- | --- | --- | --- | --- | --- | --- |
|  | 0% LD | 40% LD | 80% LD | 0% LD | 40% LD | 80% LD |
| Core genes | 2,626 | 2,012 | 432 | 1,953 | 1,314 | 357 |
| Soft core genes | 527 | 1,107 | 2,121 | 1,162 | 1,747 | 2,139 |
| Shell genes | 2,570 | 2,665 | 3,185 | 2,635 | 2,709 | 3,220 |
| Cloud genes | 14,243 | 23,320 | 34,908 | 22,461 | 27,941 | 37,014 |
| Total genes | 19,966 | 29,104 | 40,646 | 28,211 | 33,711 | 42,730 |

These findings suggest that length differences within gene families significantly influence both the size and structure of the inferred pangenome. Although sequence similarity remains central to gene clustering, allowing genes with large length disparities to cluster together can dramatically shift core and accessory gene group composition. This highlights the importance of carefully selecting parameters such as `length difference cutoff` to prevent the misgrouping of phylogenetically or functionally distinct genes. Notably, adjustments to CD-HIT `length difference cutoff` resulted in PyamilySeq drastically increasing its core gene family count to be more in line with previously reported *E. coli* pangenome studies [39, 40, 41, 42], but still lower than Roary and Panaroo’s default estimates. Core gene counts in past studies have varied widely, possibly due to differences in clustering tools, parameter settings, and core gene definitions. This variability reinforces the need for transparent, flexible, and well-documented clustering methodologies in pangenomic research, ensuring reproducibility and accuracy in gene family classification. To this end, since version 1.5.1, Panaroo now exposes the CD-HIT `length difference cutoff` parameter to address the aforementioned issues of clustering varying-length sequences together. However, this is not the default and still requires user understanding and investigation.

### 3.7 There is significant variation in single gene phylogenies

Phylogenetic trees derived from pangenomic analyses are typically constructed using two principal strategies. The first approach reconstructs phylogenies independently for each gene family and then derives a consensus or species tree from the ensemble of gene trees. The second involves concatenating the gene alignments from these independent gene trees into a single ‘super-gene’ from which a species tree is inferred. The limitations of the concatenation method, especially its sensitivity to gene tree discordance, have been recognised for over two decades [43]. Despite this, concatenation remains the default in many pangenome tools, including Roary, Panaroo, and now PyamilySeq.

To investigate the consistency of phylogenetic signal across single-gene trees, the core gene families identified by Roary, Panaroo, and PyamilySeq were analysed. As a reminder, for each tool, multiple sequence alignments were generated using MAFFT [27], and gene trees were constructed from the concatenated alignments with FastTree [30]. Pairwise RF distances were then calculated between all gene trees within each tool’s dataset to quantify their topological differences and assess overall concordance. Since Roary and Panaroo are thought to select the longest representative sequence when multiple sequences are present in a gene family, but do not report exactly which sequence was chosen, this analysis was restricted to core gene families with exactly one sequence per genome. For each tool, two versions were analysed. For Panaroo, one dataset excluded all refound sequences, while the other (Panaroo refound) included refound sequences. For Roary, results were compared between runs with the `don’t split paralogs` parameter enabled (Roary dsp) and disabled. For PyamilySeq, two datasets were generated using different `length difference cutoff` values: one with 80% and another with 0%. As shown in the all-vs-all RF distance distributions in Figure 3, the majority of single-gene trees exhibit high normalised RF distances, indicating substantial topological variation between core gene trees even within the same tool. While the differences across tools are relatively modest (see Supplementary Table 8), the lowest average normalised RF score was observed for Roary (0.892 - std:0.0695), and the highest for Panaroo (0.912 - std:0.0682). Notably, Panaroo refound includes a small set of gene trees with very low RF distances, potentially artefactual, raising concerns about the inclusion of refound sequences in phylogenetic analysis. In Supplementary Figure 10, the RF distances between each tool’s single gene trees and their corresponding concatenated tree are reported. Again, the results highlight the inconsistency between singlegene topologies and the concatenated ‘species’ tree, underscoring the need to treat such trees with caution when drawing biological conclusions.

**Figure 3:**
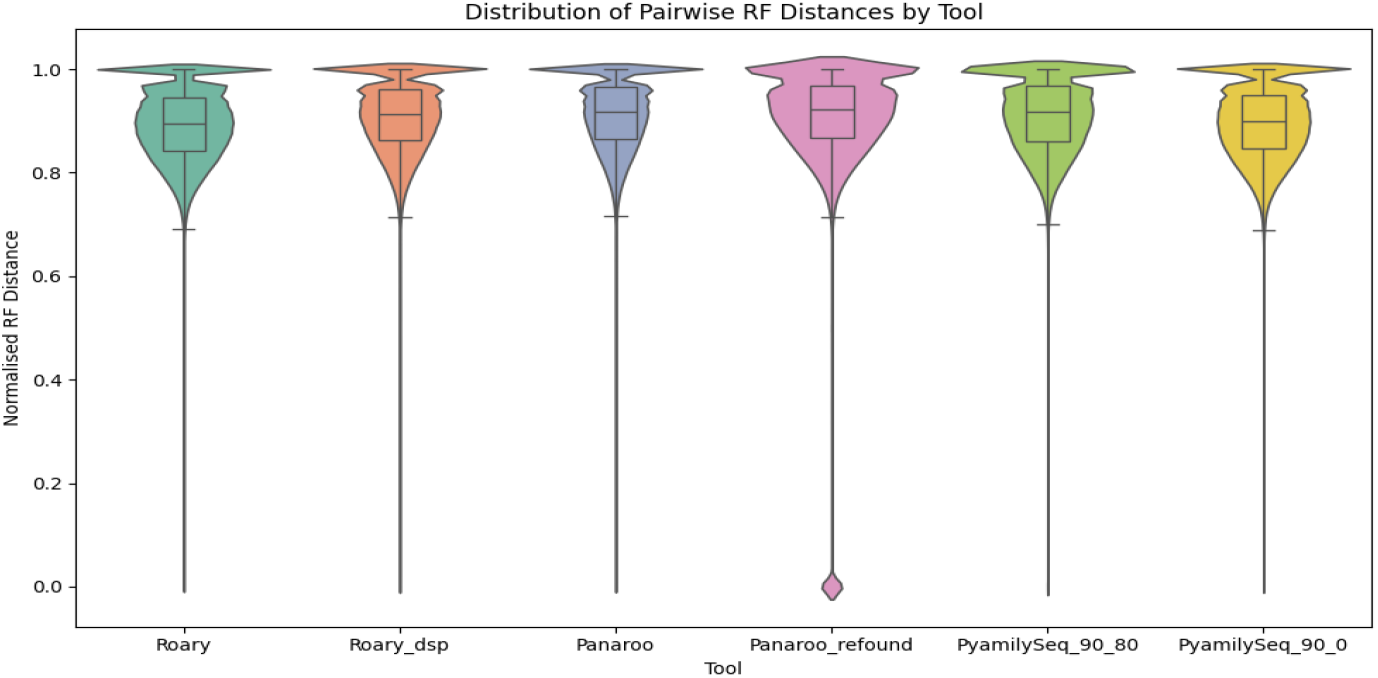
Plotted here are the normalised all-vs-all Robinson–Foulds distances computed for the core gene trees reported by the 3 different tools. Roary dsp stands for Roary performed with the `don’t split paralogs` option and Panaroo refound stands for allowing refound sequences to be processed as annotated sequences. PyamilySeq 90 80 and PyamilySeq 90 0 stand for when PyamilySeq was either run with 80% or 0% `length difference cutoff`.

### 3.8 Reclustering breaks gene clusters

Pangenome inference is only as reliable as the data it uses. The quality and completeness of both genome assemblies and gene annotations directly influence the composition of inferred gene families. In a previous study, we showed that even for well-studied bacteria such as *E. coli*, there still exist regions of its genome without annotation [13]. StORF-Reporter was then built to investigate these ‘intergenic’ regions for the types of genes contemporary annotation tools often miss [28]. Therefore, to assess whether these potentially missing gene sequences are hindering our pangenomic inference, the 74 *E. coli* genomes were re-annotated using StORF-Reporter, supplementing the original Ensembl annotations without modifying them. StORF-Reporter inserts additional CDS predictions (StORFs) into the GFF files, allowing Roary and Panaroo to process them as standard CDS sequences. Given Roary’s sensitivity to the `dont split paralogs` option, Table 6 presents the increase in core gene families following StORF-Reporter enhancement.

**Table 6:** The influence that adding additional StORF sequences has on the output of Roary (processing the 74 *E. coli* dataset). ‘**-s**’ signifies the use of the `dont split paralogs` option.

| Gene families | Roary | Roary & StORFs | Roary -s | Roary -s & StORFs |
| --- | --- | --- | --- | --- |
| Core genes | 1,023 | 1,202 | 2,554 | 2,821 |
| Soft core genes | 860 | 1,238 | 602 | 1,076 |
| Shell genes | 4,496 | 9,287 | 2,430 | 6,332 |
| Cloud genes | 31,890 | 69,651 | 16,000 | 48,021 |
| Total genes | 38,469 | 81,378 | 21,586 | 58,250 |

Panaroo, in particular, reported a significant increase, with core gene families rising from 2,824 to 3,594 (Table 7). While this might seem like a positive outcome, the results are not entirely straightforward. Ideally, the enhanced dataset should either expand existing clusters or introduce new ones, yet this is not always the case. For example, in the original clustering, group 16792 was reported with one sequence from each 74 genomes, but with the additional StORF sequences, the Ensembl annotated gene (*ENSB ru5NaBpXuypHUlL*) from genome 38344 G02, is now isolated as a singleton cluster (group 13871). This single sequence of 97 AA is the only shorter sequence in the group, where all other members are 211 AA. Strangely, despite this exclusion, much smaller StORF-derived ‘gene-copies’ remain clustered within core families. For instance, in ASM16791v2, the 53 AA StORF sequence (*StORF 62452c9dbe09d477*) is grouped with its 211 AA counterpart (*ENSB 12uI2Ow5HXCFra6*). These inconsistencies suggest internal filtering or clustering mechanisms that do not follow an immediately apparent logic, raising questions about how pangenome tools handle alternative annotations and sequence variability.

**Table 7:** Gene family distribution of the 74 *E. coli* genomes reported by Panaroo with and without the additional StORF sequences.

| Gene families | Panaroo | Panaroo & StORFs |
| --- | --- | --- |
| Core genes | 2,824 | 3,594 |
| Soft core genes | 472 | 956 |
| Shell genes | 3,040 | 6,323 |
| Cloud genes | 15,190 | 38,455 |
| Total genes | 21,526 | 49,328 |

While the exact mechanisms driving these clustering inconsistencies remain unclear, PyamilySeq provides a way to systematically track the impact of newly annotated sequences without disrupting existing clusters. By holding the original sequences and clusters in place, PyamilySeq allows us to observe where new sequences integrate, offering a controlled approach to evaluating annotation-driven changes. This resulted in several gene families that were previously classified as accessory transitioning into soft core following the incorporation of StORFs. However, the biggest changes are the entirely new core and soft-core gene families that emerged from StORF sequences alone. Supplementary Table 9 summarises the integration of the additional StORFs into existing clusters using PyamilySeq’s Partial Mode with the reclustering option. For a detailed breakdown of each gene family type, refer to Supplementary Listing 1.

### 3.9 These aren’t the genes you’re looking for

A fundamental assumption in pangenome analysis is that the ‘representative’ sequences selected for each core gene family accurately reflect the group they represent. Roary, Panaroo, and PyamilySeq all report such representative sequences, intended to serve as a centroid, the most typical or central sequence within a gene family. However, as we might now expect, this does not always hold true. Examining the *gene_presence_absence*.*csv* files reveals clear instances where representative sequences are misleading. In the Panaroo (refound-off - to ensure we are not getting sequences not in the provided annotations) analysis of the 74 *E. coli* genomes, core gene family group 15420 consists of 74 sequences, 73 of which are 564 nt / 188 AA in length. Yet, its assigned representative sequence is 1,494 nt / 498 AA, nearly three times longer. This arises because Roary and Panaroo select the longest sequence by default, rather than the most representative (however that may be defined). The 73 shorter sequences align perfectly (100% pident) with a known `membrane integrity-associated transporter subunit PqiC - WP 000759128.1` (187 AA). However, the single longer sequence aligns instead with a `paraquat-inducible protein B-STK76389.1` (497 AA), a different gene with a distinct function. This means that for group 15420, the assigned representative sequence does not match the actual gene or function of its members. This is not just a minor inconsistency; it fundamentally undermines downstream functional analysis. If a researcher used these sequences for comparative genomics, pathway reconstruction, or functional annotation, they would be working with an incorrect sequence.

Once again, this highlights the hidden pitfalls of naively using complex tools and workflows, where small, often overlooked algorithmic choices introduce systematic errors that can incorrectly influence biological interpretations. These aren’t the genes you’re looking for.

## 4 Discussion and Conclusion

Pangenomic analysis plays a critical role in understanding microbial evolution, functional diversity, and the distribution of genes across taxa, yet the accuracy and impact of its insights are fundamentally dependent on the methodologies used. This study highlights multiple sources of variation and bias in gene clustering and pangenome inference, demonstrating how parameter selection and seemingly arbitrary algorithmic decisions can significantly reshape the reported composition of core and accessory gene families. As such, widely used gene clustering and pangenome inference tools suffer from opacity, inflexibility, and methodological biases. Hard-coded thresholds, arbitrary methodological decisions, and inaccurate algorithms misgroup genes, distorting biological relationships and limiting discoveries.

Most sequence alignment or clustering tools, including BLAST (blastn for DNA, blastp for AA) and CD-HIT (cd-hit for AA and cd-hit-est for DNA), operate under the assumption that codon translation follows the universal codon table 11, applying it uniformly across genomes. However, this overlooks non-standard codons and alternative genetic codes, which, though rare, are present in certain gene families and taxa (e.g., *Mycoplasma*). Panaroo partially addresses this by clustering “concomitant genes at the DNA level”, yet its overall approach still relies heavily on homologous signal in AA sequences. To systematically investigate potential biases, PyamilySeq was used to infer gene families from both DNA and AA sequence clustering and results showed that resulting pangenomes differed.While pangenomic tools such as Roary and Panaroo have been instrumental in advancing the field, they are not immune to these methodological artefacts, and their influence on downstream analyses is often underestimated. Another key issue rehighlighted in this study is the assumption of determinism in bioinformatic tools. Many researchers operate under the assumption that running the same tool with broadly the same parameters will yield consistent results. However, this study demonstrates that factors as seemingly trivial as CPU and memory allocation, parameter floating point precision, and even output selection can all lead to drastically different gene family assignments. The lack of transparency in sequence clustering and pangenome tools has led to unexpected results. As we discovered, requesting an alignment of core gene sequences can inexplicably alter gene family composition.

More concerningly, representative sequences for core gene families are potentially selected based on length rather than biological relevance, leading to incorrect functional and phylogenetic conclusions. Such inconsistencies raise questions about the reproducibility and reliability of pangenomic studies, particularly when their conclusions influence fields such as antimicrobial resistance surveillance and pathogen detection. For example, there are tools such as Coinfinder [44] which are used in downstream studies to analyse the co-occurrence of gene families, an analysis that could be greatly affected by the issues highlighted.

Responsibility for the issues raised in this study is shared widely. Although concerns have been raised for some time [45] about whether much of the software in bioinformatics is fit-for-purpose, the development and maintenance of computational tools are often underfunded and largely driven by small teams or even solo developers, who lack the resources to support their tools in the long term. Compounding this, there is widespread misunderstanding and misuse of tools within the community. Software designed for a narrow purpose is frequently repurposed for broader applications, sometimes becoming overextended into multi-functional “Swiss army knives”, far beyond the original intent of their developers. Furthermore, papers describing novel tools are static snapshots, and documentation is often limited or quickly outdated. Many developers are domain scientists rather than trained software engineers, and even those who are still rely heavily on their user communities to report bugs or inconsistencies. This, in part, has led to issues being independently identified and attempts made to account for them, such as in a recent study by Behruznia *at al* where they used “in-house Python scripts” to rectify gene families which were “oversplit” by Panaroo [46]. Yet, with a user base that spans a broad range of expertise, it can be overwhelming, even for experienced users, to identify, understand, and communicate these problems effectively. The result is a complex and decentralised ecosystem where small errors can propagate widely, and the line between biological insight and computational artefact becomes increasingly blurred.

Additionally, while the biological significance of clustered genes is often based on sequence similarity, this study suggests that high similarity alone does not always justify their grouping. Gene birth [47], neofunctionalisation [**?**], and pseudogenisation [48] introduce complexity that is difficult, if not impossible, to account for in current clustering approaches. As demonstrated, sequences of vastly different lengths are grouped together, while near-identical sequences are excluded due to unclear rules. The field is increasingly recognising these methodological challenges. While some tools, such as Panaroo, are actively maintained and iteratively improved by dedicated developers, many tools core to sequence clustering and pangenome analysis receive limited post-publication support. Even for actively developed tools like Panaroo, correct usage requires a detailed understanding that many users lack. Problematic defaults remain unchanged, and opaque interactions between length filtering and gene grouping prevent researchers from systematically evaluating whether their specific parameter choices are biologically appropriate for their dataset.

While orthology-based tools such as OrthoMCL [49] and OrthoFinder [50] offer algorithmically sophisticated approaches to identifying gene relationships across species, they do not address the fundamental challenges we have identified in prokaryotic gene-centric pangenomic inference. These tools were primarily designed for comparative genomics across distantly related species, where the focus is on identifying orthologous genes that diverged through speciation events. In contrast, pangenome analysis of closely related prokaryotic genomes requires methods optimised for detecting recent gene duplications, horizontal gene transfer events, and high sequence similarity within species. These are biological phenomena that operate on vastly different evolutionary timescales [51]. More critically, the core algorithmic issues we have demonstrated are not unique to the tools in this study. OrthoFinder and OrthoMCL rely on similar underlying clustering heuristic tools, such as BLAST, and would face comparable challenges when applied to the high-similarity, paralog-rich landscape of prokaryotic pangenomes. In fact, the OrthoFinder paper has an entire section dedicated to the “Gene length bias in BLAST” which states that there is a “clear length dependency in the scores that are obtained… Short sequences cannot produce large bit scores or low e-values, whereas long sequences produce many hits with scores better than those for the best hits of short sequences.”.

Constructing phylogenies to represent the evolutionary relationships among genomes is a central pangenomic output. However, approaches rely on the assumption that all core genes share a common evolutionary history, an assumption that rarely, if ever, holds in prokaryotes. Gene histories can diverge due to processes such as horizontal gene transfer, recombination, or incomplete lineage sorting [52], resulting in considerable discordance between gene trees. Compounding this issue is a general lack of transparency in how core gene sequences are selected and aligned. Most tools do not clearly report which sequences are included in the final concatenated alignments, nor do they explain how multi-copy gene families are resolved. This becomes especially problematic given that the inclusion of additional sequences or paralogs does not necessarily improve phylogenetic resolution, and may in fact introduce noise or misleading signal [53]. These issues can result in trees that reflect software artefacts more than they do true biological relationships. For instance, in Panaroo’s group 21164, sequences ranging from 144 nt to 2,292 nt are clustered into a single gene family. To align these during concatenation, tools like MAFFT are forced to introduce large stretches of gaps (‘-’) to compensate for length differences. The biological relevance of such heavily gapped alignments is questionable, especially when insertion–deletion events (indels) are treated no differently from ambiguous bases such as ‘N’ by most phylogeny tools. Although this issue has been acknowledged in recent work [54], widely adopted solutions remain lacking. These limitations reinforce concerns raised in earlier studies, which showed that trees constructed from concatenated alignments often lack support from the individual gene trees they comprise. As noted by Thiergart et al., “the branches in trees made from concatenated alignments are, in general, not supported by any of their underlying individual gene trees, even though the concatenation trees tend to possess high bootstrap proportion values” [55]. This highlights the need for caution when interpreting such trees as accurate representations of species-level evolutionary histories.

To address these ongoing challenges, while not intended to replace contemporary tools, PyamilySeq has been developed to provide a flexible and transparent framework to systematically identify challenges in gene clustering and pangenomic inference, and thus, guide the development of more transparent methodologies. By allowing researchers to explore the impact of clustering parameters, PyamilySeq provides an unparalleled level of transparency compared to existing tools. It enables iterative reclustering, allowing newly annotated sequences to be incorporated without disrupting existing clusters, but also provides the ability to track the disrupted clusters and sequences. It is presented with several additional tools that can help the exploration of previous clustering or pangenome results to help researchers understand any potential problems or oddities that may exist. PyamilySeq complements contemporary and future approaches by enabling the exposure of their algorithmic decisions and consequences and underscores the need for greater user awareness of the impact of methodological choices, reinforcing the importance of parameter sensitivity analysis in sequence clustering and pangenomics. Future efforts should focus on integrating biologically informed clustering techniques that account for evolutionary processes beyond sequence similarity alone. By critically evaluating the tools that define pangenomes, we can move toward more accurate, reproducible, and biologically meaningful insights.

## 5 Data availability

The genomic data used are listed in Supplementary Tables 1 and 2 and are available at Ensembl Bacteria. Data files created are available on figshare 10.6084/m9.figshare.29424626.

## 7 Author contributions statement

N.J.D undertook all development of PyamilySeq, investigated the clustering and pangenome tools studied and wrote the manuscript.

## 8 Acknowledgments

N.J.D. acknowledges Prof. Michael Surette and Dr. Katie Lawther for their invaluable advice, which significantly improved the manuscript. Additional thanks to Dr. Amanda Clare and Dr. Fiona Whelan for their thoughtful comments and continued support of the project, and also to the 2025 EMBO Practical Course (Computational Molecular Evolution) organisers and participants for their constructive discussions and encouragement.

