## Supplementary File 1 for "PyamilySeq: Exposing the fragility of conventional gene (re)clustering and prokaryotic pangenomic inference methods" (from another version of this preprint: version 2, 2025-06-27)

Nicholas J. Dimonaco [1,2*]

[1]Institute for Global Food Security, School of Biological Sciences, Queen's University Belfast, Belfast, BT9 5DL, UK
[2]Department of Computer Science, Aberystwyth University, Aberystwyth, SY23 3DB, Wales, UK

### 1 Supplementary Figures

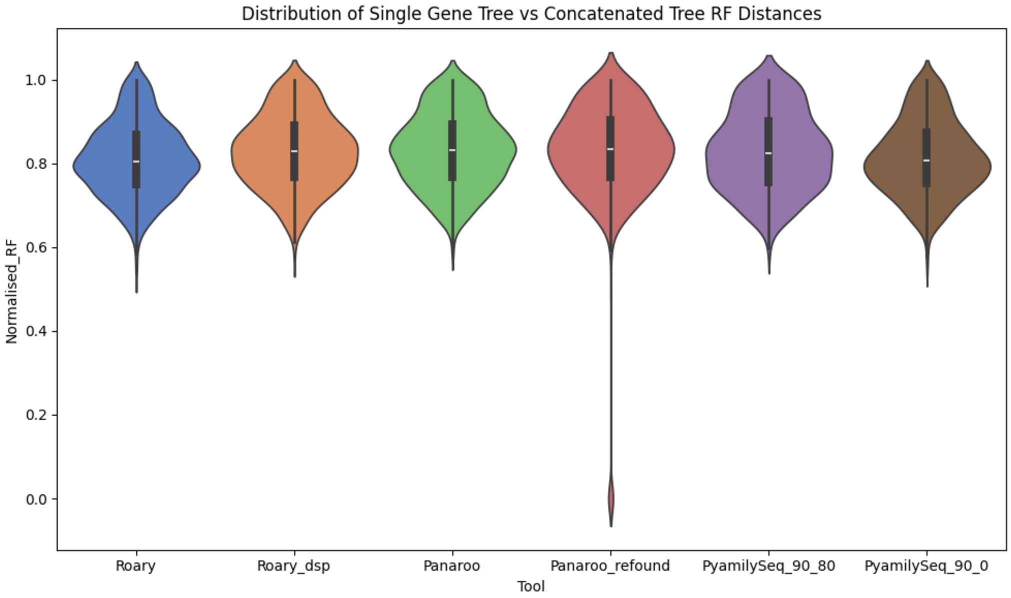


Supplementary Figure 1: The normalised Robinson–Foulds distances computed for each tools set of core single gene trees against the concatenated gene trees. Roary_dsp stands for Roary performed with the `don't split paralogs` option and Panaroo_refound stands for allowing refound sequences to be processed as annotated sequences.

#### 2 Supplementary Tables

| COG Group | PyamilySeq [%] | Roary [%] | Panaroo [%] |
|---|---|---|---|
| ISP | 97 [**26.15**] | 250 [**23.61**] | 548 [**18.72**] |
| CPS | 62 [**16.71**] | 185 [**17.47**] | 586 [**20.02**] |
| MET | 142 [**38.27**] | 410 [**38.71**] | 1,298 [**44.34**] |
| PC | 70 [**18.86**] | 214 [**20.21**] | 495 [**16.91**] |

Supplementary Table 1: Reported here are the broad functional COG categories in raw numbers and proportions. The COG group acronyms are as follows: ISP - INFORMATION STORAGE AND PROCESSING, CPS - CELLULAR PROCESSES AND SIGNALING, MET - METABOLISM and PC - POORLY CHARACTERIZED.

| Groups | 1 CPU | | | 8 CPU | | |
|---|---|---|---|---|---|---|
| | 8,000 MB | 4,000 MB | 800 MB | 8,000 MB | 4,000 MB | 800 MB |
| Core genes: | 1952 | 1952 | 1953 | 1952 | 1952 | 1953 |
| Soft core genes: | 1160 | 1160 | 1160 | 1160 | 1160 | 1160 |
| Shell genes: | 2644 | 2644 | 2640 | 2644 | 2644 | 2640 |
| Cloud genes: | 22455 | 22455 | 22458 | 22455 | 22455 | 22458 |
| Total: | 28211 | 28211 | 28211 | 28211 | 28211 | 28211 |

Supplementary Table 2: The slightly different clustering results we get for the number of core genes due to the different CPU allocation. Memory allocation does not seem to make a difference, at least at the.....

| CD-HIT initialisations | 1 CPU | | | 8 CPU | | |
|---|---|---|---|---|---|---|
| | 8,000 MB | 4,000 MB | 800 MB | 8,000 MB | 4,000 MB | 800 MB |
| Buffer (**MB**) | 22 | 22 | 22 | 176 | 176 | 176 |
| Table (**MB**) | 71 | 71 | 71 | 143 | 143 | 143 |
| Total (**MB**) | 283 | 283 | 283 | 509 | 509 | 509 |
| Representatives | 40,000,000 | 40,000,000 | 1,054,982 | 40,000,000 | 40,000,000 | 593,077 |
| Word Count Entries | 964,607,788 | 464,607,788 | 64,607,788 | 936,320,494 | 436,320,494 | 36,320,494 |

Supplementary Table 3: The different initialisation setups that CD-HIT reports based on different CPU and memory allocations.

| Tool | Normalised Robinson-Foulds | Standard Deviation |
|---|---|---|
| Roary | 0.8920 | 0.0695 |
| Roary_dsp | 0.9080 | 0.0683 |
| Panroo | 0.9120 | 0.0682 |
| Panroo_refound | 0.8937 | 0.1597 |
| PyamilySeq_90_80 | 0.9108 | 0.0732 |
| PyamilySeq_90_0 | 0.8956 | 0.0708 |

Supplementary Table 4: Reported here are the normalised all-against-all Robinson-Foulds distances computed from the single gene trees from each of the tools studied. Roary_dsp stands for Roary performed with the `don't split paralogs` option and Panaroo_refound stands for allowing refound sequences to be processed as annotated sequences.

| **Genome** | **Number of Contigs** | **Number of CDSs** |
|---|---|---|
| Escherichia_coli_110957_gca_000485615.ASM48561v1 | 241 | 4931 |
| Escherichia_coli_1_392_07_s4_c1_gca_000713675.ASM71367v1 | 412 | 5565 |
| Escherichia_coli_5_366_08_s1_c1_gca_000714255.ASM71425v1 | 123 | 5145 |
| Escherichia_coli_b185_gca_000163175.ASM16317v1 | 44 | 4675 |
| Escherichia_coli_536_gca_000013305.ASM1330v1 | 1 | 4540 |
| Escherichia_coli_2362_75_gca_000183005.ASM18300v1 | 83 | 4965 |
| Escherichia_coli_4_0522_gca_000194335.ASM19433v2 | 109 | 5780 |
| Escherichia_coli_2_177_06_s3_c2_gca_000704005.ASM70400v1 | 181 | 5040 |
| Escherichia_coli_113303_gca_000485655.ASM48565v1 | 101 | 4734 |
| Escherichia_coli_113290_gca_000485635.ASM48563v1 | 189 | 4952 |

Supplementary Table 5: The 10 *Escherichia coli* genomes used in ‘Case Study: Pangenomic Analysis of 10 *Escherichia coli* Genomes’ as reported from Ensembl Bacteria release 59.

| Genome | Number of Contigs | Number of CDSs |
|---|---|---|
| Escherichia_coli_o25b_h4_gca_005671235.ASM567123v1 | 3 | 4966 |
| Escherichia_coli_o25b_h4_gca_005671255.ASM567125v1 | 4 | 5117 |
| Escherichia_coli_o25b_h4_gca_005673435.ASM567343v1 | 5 | 5033 |
| Escherichia_coli_o55_h7_str_cb9615_gca_000025165.ASM2516v1 | 2 | 5146 |
| Escherichia_coli_o7_k1_str_ce10_gca_000227625.ASM22762v1 | 5 | 5140 |
| Escherichia_coli_sms_3_5_gca_000019645.ASM1964v1 | 5 | 4819 |
| Escherichia_coli_umn026_gca_000026325.ASM2632v2 | 3 | 5015 |
| Escherichia_coli_uti89_gca_000013265.ASM1326v1 | 2 | 4837 |
| Escherichia_coli_w_gca_000184185.ASM18418v1 | 3 | 4703 |
| Escherichia_coli_nccp15648_gca_003433275.ASM343327v1 | 4 | 5357 |
| Escherichia_coli_o103_h2_str_12009_gca_000010745.ASM1074v1 | 2 | 5334 |
| Escherichia_coli_o111_nm_gca_005037805.ASM503780v2 | 1 | 4917 |
| Escherichia_coli_o127_h6_gca_900149915.EPEC-E2348_69-V2 | 1 | 4637 |
| Escherichia_coli_o127_h6_str_e2348_69_gca_000026545.ASM2654v1 | 3 | 4872 |
| Escherichia_coli_o157_gca_002208865.ASM220886v2 | 3 | 5498 |
| Escherichia_coli_o157_h7_str_edl933_gca_000006665.ASM666v1 | 2 | 5416 |
| Escherichia_coli_o25b_h4_gca_005670925.ASM567092v1 | 2 | 5171 |
| Escherichia_coli_o25b_h4_gca_005671055.ASM567105v1 | 3 | 4937 |
| Escherichia_coli_gca_900449115.38828_C01 | 4 | 6403 |
| Escherichia_coli_gca_900449225.38891_G01 | 5 | 5180 |
| Escherichia_coli_gca_900449295.39038_F01 | 3 | 6105 |
| Escherichia_coli_gca_900449435.38891_F02 | 4 | 5142 |
| Escherichia_coli_gca_900449455.38891_E02 | 4 | 5217 |
| Escherichia_coli_gca_900449865.39282_C02 | 5 | 7142 |
| Escherichia_coli_gca_900449985.40363_C01 | 5 | 5101 |
| Escherichia_coli_gca_900450185.40105_F02 | 2 | 5436 |
| Escherichia_coli_gca_900520285.ASM90052028v1 | 2 | 4751 |
| Escherichia_coli_gca_900607665.ecoli009 | 5 | 5687 |
| Escherichia_coli_gca_001900535.ASM190053v1 | 3 | 5427 |
| Escherichia_coli_gca_003769125.ASM376912v2 | 3 | 4985 |
| Escherichia_coli_gca_006351885.ASM635188v1 | 5 | 5337 |
| Escherichia_coli_gca_900449095.38891_C01 | 4 | 6494 |
| Escherichia_coli_gca_901733115.Escherichia_coli_MAD | 4 | 4585 |
| Escherichia_coli_o25b_h4_gca_005671075.ASM567107v1 | 2 | 4816 |
| Escherichia_coli_gca_006364695.ASM636469v1 | 4 | 5107 |
| Escherichia_coli_gca_900184875.Unitig | 4 | 4878 |
| Escherichia_coli_gca_900447915.44087_F02 | 3 | 4897 |
| Escherichia_coli_gca_900448265.46338_F02 | 5 | 5212 |
| Escherichia_coli_gca_900448335.50624_F01 | 5 | 5122 |
| Escherichia_coli_gca_900448615.38344_G02 | 2 | 7321 |
| Escherichia_coli_gca_900448835.38389_B01 | 4 | 5222 |
| Escherichia_coli_gca_900448935.38670_B01 | 3 | 5138 |
| Escherichia_coli_gca_900448955.38656_D02 | 4 | 6283 |
| Escherichia_coli_gca_900448985.38490_E02 | 4 | 5584 |
| Escherichia_coli_gca_003790525.ASM379052v1 | 5 | 5914 |
| Escherichia_coli_gca_003991155.ASM399115v1 | 5 | 5304 |
| Escherichia_coli_gca_004011015.ASM401101v1 | 3 | 5202 |
| Escherichia_coli_gca_004114395.ASM411439v1 | 4 | 5057 |
| Escherichia_coli_gca_004358365.ASM435836v1 | 3 | 5666 |
| Escherichia_coli_gca_004564175.ASM456417v1 | 3 | 4993 |
| Escherichia_coli_gca_005890115.ASM589011v1 | 1 | 4474 |
| Escherichia_coli_gca_005954725.ASM595472v1 | 1 | 4484 |
| Escherichia_coli_gca_006337025.ASM633702v1 | 3 | 4933 |
| Escherichia_coli_gca_001902655.ASM190265v1 | 3 | 5297 |
| Escherichia_coli_gca_002011985.ASM201198v1 | 4 | 5100 |
| Escherichia_coli_gca_002156845.ASM215684v1 | 3 | 5513 |
| Escherichia_coli_gca_002164595.ASM216459v2 | 5 | 5052 |
| Escherichia_coli_gca_002554485.ASM255448v1 | 5 | 5240 |
| Escherichia_coli_gca_002846135.ASM284613v1 | 3 | 5418 |
| Escherichia_coli_gca_002860085.ASM286008v1 | 5 | 5176 |
| Escherichia_coli_gca_003122105.ASM312210v1 | 4 | 4930 |
| Escherichia_coli_gca_003571785.ASM357178v1 | 5 | 5129 |
| Escherichia_coli_53638_gca_000167915.ASM16791v2 | 2 | 4924 |
| Escherichia_coli_536_gca_000013305.ASM1330v1 | 1 | 4540 |
| Escherichia_coli_55989_gca_000026245.ASM2624v1 | 1 | 4841 |
| Escherichia_coli_apec_o1_gca_000014845.ASM1484v1 | 3 | 5254 |
| Escherichia_coli_cft073_gca_000007445.ASM744v1 | 1 | 4900 |
| Escherichia_coli_chi7122_gca_000307205.ASM30720v1 | 1 | 4496 |
| Escherichia_coli_gca_000801185.ASM80118v2 | 2 | 4679 |
| Escherichia_coli_gca_001420955.ASM142095v1 | 4 | 5249 |
| Escherichia_coli_gca_001515725.ASM151572v1 | 1 | 4557 |
| Escherichia_coli_gca_001612475.ASM161247v1 | 2 | 5434 |
| Escherichia_coli_gca_001677475.ASM167747v1 | 2 | 5477 |
| Escherichia_coli_gca_001721525.ASM172152v1 | 5 | 5307 |

Supplementary Table 6: The 74 *Escherichia coli* genomes used in 'Case Study: Pangenomic Analysis of 74 *Escherichia coli* Genomes' as reported from Ensembl Bacteria release 59.

### 3 Supplementary Listings

Listing 1: CD-HIT initialisation with PyamilySeq Full mode default parameters (-T 8 and -M 4000)

```
total seq: 50325
longest and shortest : 6925 and 29
Total letters: 15353668
Sequences have been sorted
Approximated minimal memory consumption:
Sequence        : 24M
Buffer          : 8 X 12M = 100M
Table           : 2 X 66M = 132M
Miscellaneous   : 0M
Total           : 258M
Table limit with the given memory limit:
Max number of representatives: 4000000
Max number of word counting entries: 467728268
```

Listing 2: CD-HIT initialisation using 8 CPU threads but CD-HITs default memory (-T 8 and -M 800)

```
total seq: 50325
longest and shortest : 6925 and 29
Total letters: 15353668
Sequences have been sorted
Approximated minimal memory consumption:
Sequence        : 24M
Buffer          : 8 X 12M = 100M
Table           : 2 X 66M = 132M
Miscellaneous   : 0M
Total           : 258M
Table limit with the given memory limit:
Max number of representatives: 1090357
Max number of word counting entries: 67728268
```

Listing 3: CD-HIT initialisation using CD-HITs default parameters (-T 1 -M 800)

```
total seq: 50325
longest and shortest : 6925 and 29
Total letters: 15353668
Sequences have been sorted
Approximated minimal memory consumption:
Sequence        : 24M
Buffer          : 1 X 12M = 12M
Table           : 1 X 66M = 66M
Miscellaneous   : 0M
Total           : 104M
Table limit with the given memory limit:
Max number of representatives: 1400522
Max number of word counting entries: 86994327
```

### 4 Processing

#### 4.1 Input handling with Seq-Combiner

A major hurdle in bioinformatic analysis is file format compatibility. As highlighted in my previous work on the ORForise genome annotation comparison platform [1], each annotation tool delivers its own output format, and even tools pertaining to use the General Feature Format (GFF), often have tool-specific interpretations. Tools that process genome annotation output, frequently struggle with these varying formats, requiring users to modify annotations to meet specific input requirements. For example, Roary and Panaroo require their input to be Prokka-formatted [2] GFF files.

To address these compatibility issues, the PyamilySeq toolkit includes `Seq-Combiner`, which generates compatibility-assured input files for PyamilySeq. Seq-Combiner takes a directory of matching genome FASTA and GFF files or GFF files with '##FASTA - Seq' appended to the bottom and prepends the file name to each gene ID. Additionally, to support the use of other sequence clustering techniques, Seq-Combiner can process pre-extracted FASTA files and their sequences into the required naming convention, ensuring they can be seamlessly used within PyamilySeq. This flexibility ensures compatibility with diverse input data formats generated by various genome annotation and alignment tools.

An example of the sequence renaming process, which uses the names of the GFF/FASTA files, is shown below:

Listing 4: Seq-Combiner Sequence renaming.

```
GFF/FASTA Filename: Escherichia_coli..[.gff/.fasta]
GeneID: E1CxwDWD5uDlWpi

>Escherichia_coli..|E1CxwDWD5uDlWpi
>Escherichia - [GenusID]
>Escherichia_coli.. - [SpeciesID/GenomeID]
>Escherichia_coli..|E1CxwDWD5uDlWpi - [GeneID]
```

#### 4.2 Cluster summaries and investigation

The Cluster-Summary tool provides a detailed overview of CD-HIT clustering results, helping researchers evaluate and optimise clustering parameters. By analysing gene clustering patterns across genomes, it enables users to observe how parameter selection can affect results, such as the number of gene families, the distribution of core and accessory genes, and the presence of multicopy gene clusters.

Cluster-Summary processes the CD-HIT .clstr files and extracts key statistics for each cluster. The tool calculates and outputs comprehensive summaries for each cluster, including: The number of sequences in the cluster, The number of Genomes that the sequences are present in, Average sequence length, Length range (minimum to maximum), Average percentage identity, Identity range (minimum to maximum), The number of genomes with $\geq$1 gene and the percentage of genomes with $\geq$ gene.

A key feature of Cluster-Summary is its ability to assess multicopy genes, identifying clusters where genomes contain multiple representatives. It categorises these by predefined percentage ranges (e.g., 20–40%, 40–60%), providing insights into gene duplication and paralogy. This categorisation helps determine whether clustering parameters accurately reflect dataset biology, informing decisions on core, accessory, or duplicated gene families.

Additionally, PyamilySeq includes Group-Splitter, which applies Levenshtein distance [3] to refine gene clusters. This tool was used in this study to better decipher the impacts of clustering limitations and sequence divergence in the reported gene families.
